# Non-random segregation of sister chromosomes by *Escherichia coli* MukBEF axial cores

**DOI:** 10.1101/2020.07.23.217539

**Authors:** Jarno Mäkelä, Stephan Uphoff, David J. Sherratt

## Abstract

The *Escherichia coli* structural maintenance of chromosomes complex, MukBEF, forms axial cores to chromosomes that determine their spatio-temporal organization. Here, we show that axial cores direct chromosome arms to opposite poles and generate the translational symmetry between newly replicated sister chromosomes. MatP, a replication terminus (*ter*) binding protein prevents chromosome rotation around the longitudinal cell axis by displacing MukBEF from *ter*, thereby maintaining the linear shape of axial cores. During DNA replication, MukBEF action directs lagging strands towards the cell center, marked by accumulation of DNA-bound β_2_-clamps in the wake of replisomes, in a process necessary for the translational symmetry of sister chromosomes. Finally, the ancestral (‘immortal’) template DNA strand, propagated from previous generations, is preferentially inherited by the cell forming at the old pole, dependent on MukBEF-MatP. The work demonstrates how chromosome organization-segregation can foster non-random inheritance of genetic material and provides a framework for understanding how chromosome conformation and dynamics shape subcellular organization.

## Introduction

Faithful chromosome propagation and inheritance underpin all replicative life. Organisms have evolved a vast range of mechanisms to ensure timely replication and segregation of genetic material. Despite this diversity, highly conserved Structural Maintenance of Chromosomes (SMC) complexes play a central role in the organization of chromosomes in all domains of life. Eukaryotic cells orchestrate replication and segregation in discrete stages where newly replicated sister chromosomes are first individualized by condensin and held together by cohesin before being pulled apart by action of the mitotic spindle and cleavage of cohesion (reviewed in (Uhlmann, 2016)). In contrast, in prokaryotes, chromosome replication and segregation are generally not temporally separated and occur progressively (Kuzminov, 2014). Because divergent species have evolved different solutions to the same problem, understanding the contributions of different mechanisms and physical constraints underlying robust chromosome segregation remains a challenge (Badrinarayanan et al., 2015; Surovtsev and Jacobs-Wagner, 2018; Wang et al., 2013).

Genetic studies have identified two major classes of proteins implicated in chromosome segregation in bacteria. First, structural maintenance of chromosomes (SMC) complexes, MukBEF, MksBEF and Smc-ScpAB, were initially identified in a screen for *Escherichia coli* mutants that generated anucleate cells as a consequence of a failure to segregate newly replicated chromosomes to daughter cells (Hiraga et al., 1989; Nolivos and Sherratt, 2014). Second, studies of low copy plasmid stability identified ParAB*S* systems, which subsequently were shown to have roles in chromosome segregation in many organisms (Surovtsev and Jacobs-Wagner, 2018). While many bacteria encode one or both of these systems, some, for example *Pseudomonas aeruginosa*, encode two different SMCs and a ParAB*S* system (Petrushenko et al., 2011; Vallet-Gely and Boccard, 2013). Nevertheless, deletion of SMC or ParAB proteins has frequently modest if any consequences for chromosome segregation. Consistent with this, it has been proposed that large bacterial chromosomes can utilise repelling entropic effects to facilitate separation of bacterial chromosomes (Jun and Mulder, 2006), unlike much smaller low copy number plasmids that require a functional ParAB*S* system for faithful segregation (Surovtsev and Jacobs-Wagner, 2018). Whatever roles entropic forces may play, studies in diverse bacterial species have demonstrated that chromosomal loci are not positioned randomly in cells (Fogel and Waldor, 2005; Umbarger et al., 2011; Vallet-Gely and Boccard, 2013; Wang et al., 2005, 2006, 2014), and that in *E. coli*, MukBEF complexes play an important role in correct positioning of replication origins and other loci by forming an axial core to the chromosome (Danilova et al., 2007; Mäkelä and Sherratt, 2020). Absence of MukBEF leads to formation of anucleate cells during growth and loss of viability at temperatures higher than 22 °C in rich media (Danilova et al., 2007; Niki and Jaffe, 1991).

In new-born *E. coli* cells with non-overlapping replication cycles, origins of replication (*oriC*) are positioned close to the cell center, and the left and right chromosome arms are linearly organized in separate cell halves. Chromosome replication-segregation leads to generation of daughter cells with a chromosome organization identical to their mother cell. Most cell adopt a *left-oriC-right-left-oriC-right* (*L-R-L-R*) translational symmetry prior to division (Wang et al., 2006), which requires that either the leading or lagging strand templates are symmetrically segregated to the cell poles (Toro and Shapiro, 2010; Wang et al., 2005). In agreement, an elegant chromosome degradation experiment showed that the leading strand templates are segregated towards the cell poles in most cells (White et al., 2008). In theory, cells could also control the fate of the old template strand by non-random segregation, designating the destination for each strand. During each replication cycle, there is a risk of the new strand not faithfully copying the information from the template strand. ‘Immortal’ (or ancestral) strand retention was originally proposed as a strategy to maintain DNA purity in stem cells while the copied strands, potentially carrying mutations from replication, were segregated to non-stem cell progeny (Cairns, 1975). Whether this strategy is actually utilized by stem cells remains controversial (Lansdorp, 2007; Rando, 2007; Wakeman et al., 2012). Ancestral strand segregation has also been tested in *C. crescentus* (Marczynski et al., 1990; Osley and Newton, 1974) and *B. subtilis* (Errington and Wake, 1991), however, none of these studies showed any segregational strand preference between daughter cells.

We lack a mechanistic understanding of how chromosome conformation and orientation is maintained inside a bacterial cell. It also remains unknown how progressive chromosome segregation facilitates non-random sister chromosome inheritance in an otherwise apparently symmetrical organism. Here, we address these questions in *E. coli* utilizing microfluidics culturing devices combined with time-lapse imaging, high-throughput microscopy and quantitative analysis. We first demonstrate that in the absence of MukBEF, anucleate cells arise predominantly from the mother cell’s new pole as a consequence of the failure to segregate newly replicated origins in a timely fashion. We show that nascent lagging strands and their templates are directed towards cell centers, a process that is required for the observed translational *L-R-L-R* segregational symmetry; and which is perturbed in the absence of MukBEF. Furthermore, we show directly that the ancestral DNA strand, inherited from previous generations, is preferentially segregated to the old cell pole dependent on both MukBEF and its partner MatP. Lack of MatP does not perturb translational *L-R-L-R* symmetry; rather it leads to flipping of chromosome orientation around the longitudinal cell axis during a cell cycle, consistent with the observed loss of ancestral strand retention at old poles. Taken together, the results explain how MukBEF axial cores and their MatP-driven depletion from the *ter* region, lead to asymmetric strand and chromosome segregation. The possible functional and evolutionary consequences of this are explored.

## Results

### In the absence of MukBEF, anucleate cells arise from the newer mother cell pole

To understand how anucleate *E. coli* cells form in the absence of MukBEF, we followed successive cell cycles of *ΔmukB* cells with *oriC* and *ter* (*ori1* and *ter3*, respectively) regions fluorescently labeled by FROS markers. We used a ‘mother machine’ microfluidics device (Uphoff, 2018; Wang et al., 2010) to follow thousands of cell generations and identify changes in chromosome organization that correlate with chromosome mis-segregation. Under these conditions, 15.7 ± 0.4% (±SD) of divisions led to the formation of an anucleate daughter cell (Fig. 1A, Fig. S1).

**Fig. 1.**
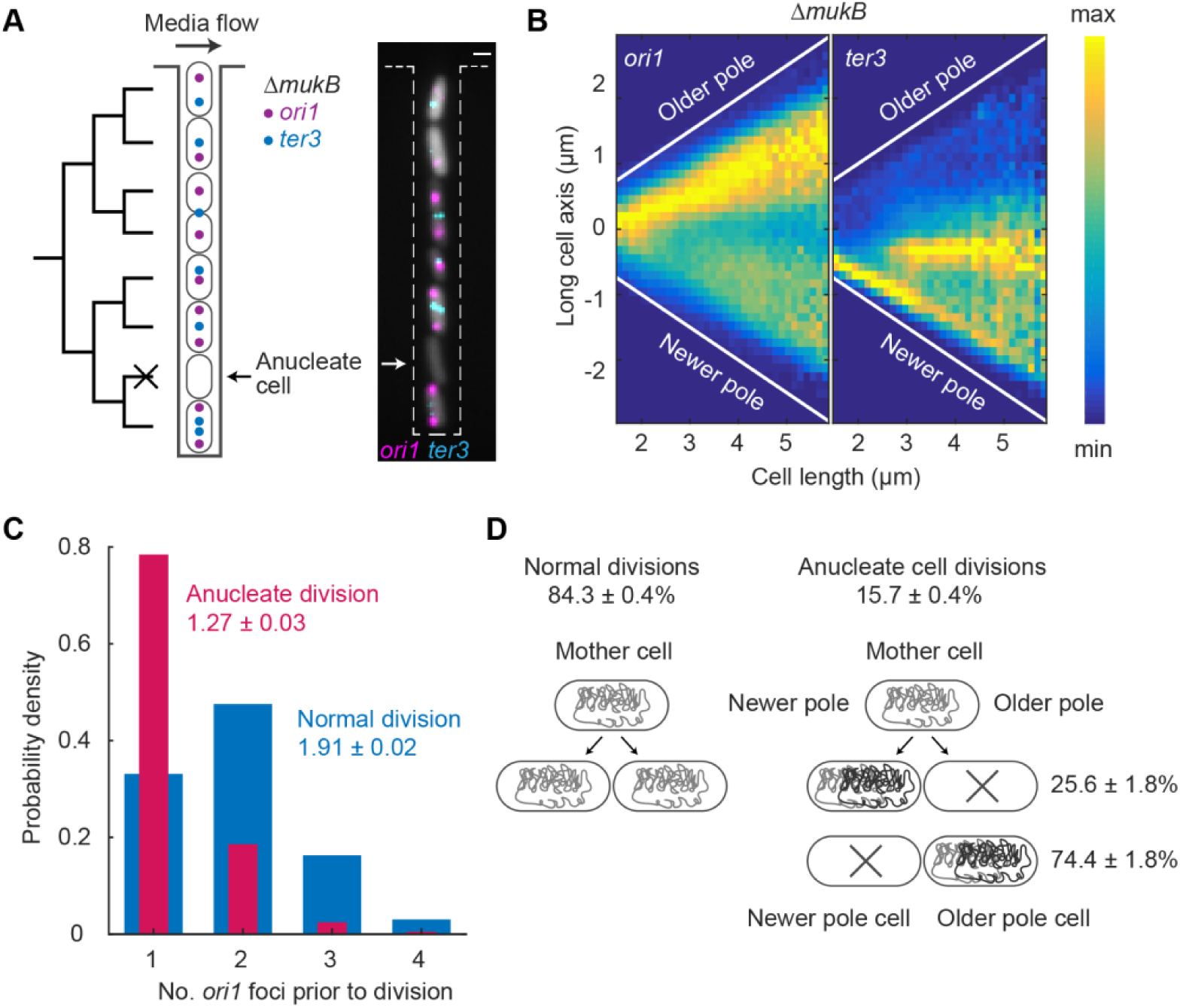
Anucleate cell formation in absence of MukB is biased towards mother cell newer poles. (**A**) Schematic of mother machine microfluidics device and representative cells in a channel. *ΔmukB* cells contain *ori1* and *ter3* FROS markers and a segmentation marker (grey). A non-growing anucleate cell lacking FROS markers is indicated. Scale bar: 1 μm. (**B**) *ori1* and *ter3* localization as a function of cell length in *ΔmukB* cells. Sample numbers with different cell lengths are normalized. 221057 cells. (**C**) Number of *ori1* foci prior to anucleate (one of the daughters is anucleate; 2444 cells) and normal cell division (10468 cells). Two-sample t-test between mean *ori1* numbers prior to anucleate and normal division p-value < 10^−5^. (**D**) Percentage of anucleate cells forming at a mother cell’s old and newer poles (2269 divisions). Percentage of anucleate cell divisions also shown (14392 divisions). All data from 3 repeats.

In *ΔmukB* cells *ori1* was of often mis-localized towards the old pole at birth, rather than at midcell (Fig. 1B), as previously observed for wild type (WT) cells (Wang et al., 2006). As the cell cycle progressed, *ori1* localized preferentially towards the old cell pole, with newly replicated sister *ori1* loci frequently remaining in close proximity. In contrast, *ter3* migrated from the new-born cell pole to midcell similar to what has been previously reported (Fig. 1B)(Wang et al., 2005). ∼80% of anucleate cells were generated when duplicated *ori1* loci in mother cells remained together in the region of the old pole prior to cell division (Fig. 1C). In contrast, in ∼70% of mother cells where chromosome segregation was faithful, *ori1* loci were visible as separate foci (Fig. 1C). Delayed separation of newly replicated *ori1* loci could be a consequence of delayed decatenation, since the decatenase TopoIV, is no longer recruited by MukBEF to *oriC*-proximal regions (Zawadzki et al., 2015). Indeed, modest over-expression of TopoIV leads to a reduction in cohesion time of newly replicated *oriC* from ∼14 min to ∼5 min (Wang et al., 2008). Delayed *ori1* decatenation of *ΔmukB* cells might explain non-viability under fast growth conditions, while slow growth conditions allow sufficient time for chromosome decatenation and segregation in most cells. In anucleate cell divisions, daughter cells that inherited two chromosomes divided normally after a modest increase in generation time (Fig. S1). However, the probability of these cells forming an anucleate cell in subsequent division was 9.1 ± 2% (±SD), significantly lower than for cells born with a single chromosome.

Prior to anucleate cell formation, mother cells divided nearly symmetrically (2.1 ± 0.2 μm and 2.4 ± 0.2 μm, respectively (±SD); two-sample t-test p-value 0.17), with the divisome being placed close to midcell. While the average anucleate cell length at birth did not significantly differ from that of the sister, the growing sister was systematically longer than the anucleate sister at birth (Fig. S1), the bias for the longer growing sister increasing with mother cell division size. Importantly, we showed that anucleate cells formed preferentially at the newer mother cell pole (74.4 ± 1.8% (±SD), Fig. 1D). Therefore, anucleate cell formation is associated with the nucleoid being preferentially retained at the old pole of *ΔmukB* mother cells, while in the case of WT cells the nucleoid is localized closer to the newer pole of the dividing cell (Fisher et al., 2013). We conclude that mis-localization of *ori1* towards the old pole, accompanied by delayed segregation of newly replicated *ori1* loci, directs the formation of anucleate cells to the mother cell’s new pole.

### MukBEF and MatP direct left-oriC-right chromosome organization

Next we investigated how MukBEF orchestrates chromosome organization and segregation in growing cells. MukBEF and MatP have been proposed to be major factors in dictating *left-oriC-right* (*L*-*R*) chromosome organization in *E. coli* (Mäkelä and Sherratt, 2020; Wang et al., 2006). MukBEF complexes form axial cores that linearly organize the chromosome outside the 800 kb *ter* region, from where MatP bound to *matS* sites displaces MukBEF (Fig. 2A)(Mäkelä and Sherratt, 2020). Complete axial cores were most easily visualized in cells in which MukBEF occupancy on the chromosome was increased ∼3.5-fold, while cells with WT MukBEF abundance on chromosomes exhibited more granular structures (Mäkelä and Sherratt, 2020). To characterize how MukBEF axial cores influence chromosome organization during the cell cycle, we used strains that allowed us to test the requirements for *left* and *right* chromosome arm organization in relation to *oriC* and *ter* in MatP^+^ and *ΔmatP* cells with WT levels of MukBEF, and in *ΔmukB* cells (Fig. 2). The *left* and *right* chromosome arms were labeled at *L3* and *R3* (−128° and 122° from *oriC*, respectively) with FROS markers, as were *ori1* and *ter3* loci (Fig. 2A, B).

**Fig. 2.**
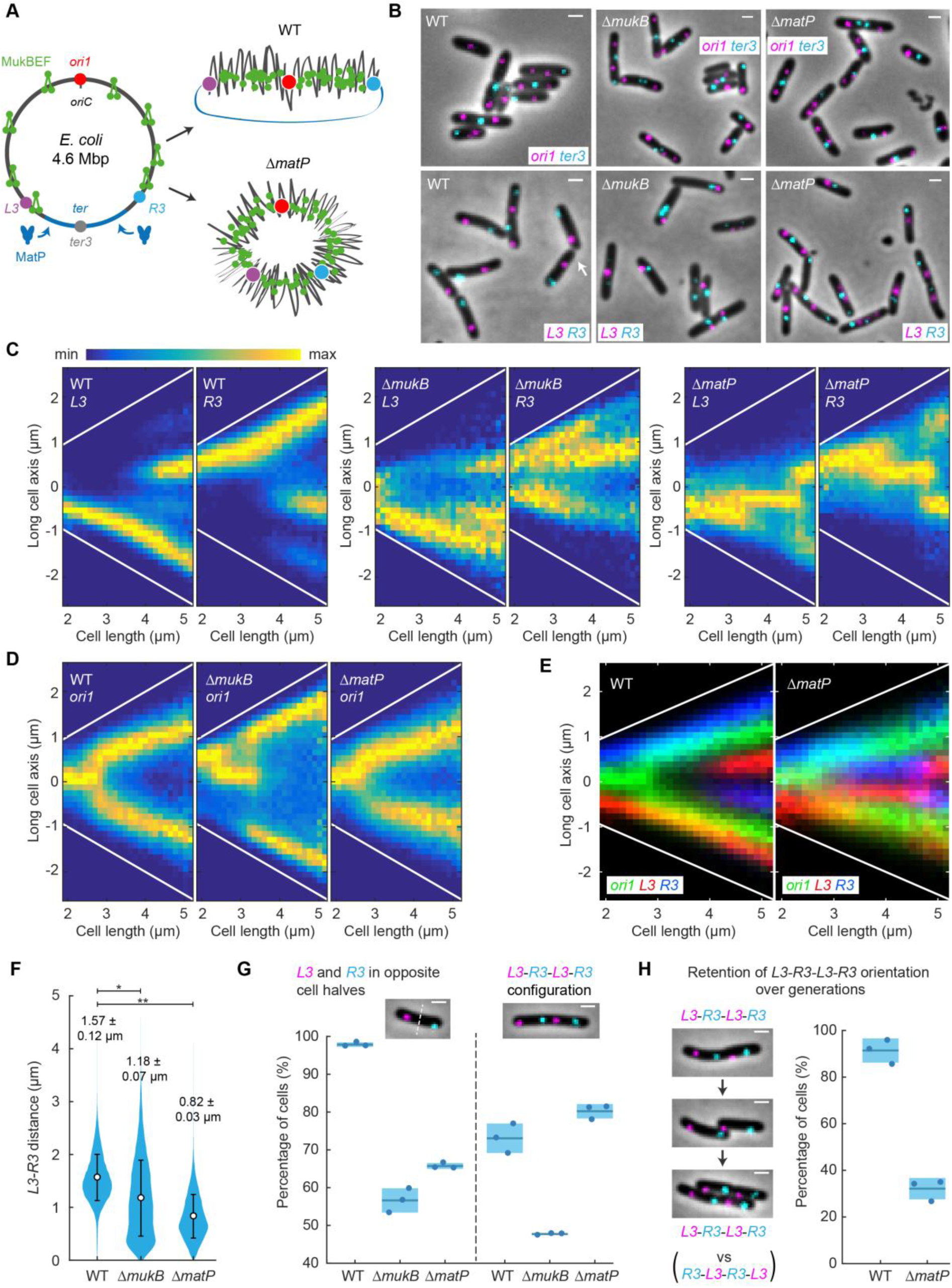
MukBEF and MatP action generates and retains *left-oriC-right E. coli* chromosome organization. (**A**) *E. coli* chromosome circular map with *ori1, ter3, L3*, and *R3* loci illustrates uniform MukBEF occupancy except for 800 kbp *ter*, from which *matS* bound MatP displaces MukBEF. A folded chromosome conformation by linear MukBEF axial cores is shown with and without MatP (for more details see (Mäkelä and Sherratt, 2020). (**B**) Representative images of WT, *ΔmukB* and *ΔmatP* cells with *ori1* and *ter3*, or *L3* and *R3* FROS markers. Note an atypical *R3-L3-L3-R3* configuration in WT (white arrow) in comparison to standard *L3-R3-L3-R3*. Scale bars: 1 μm. (**C**) *L3* and *R3* localizations and (**D**) *ori1* localizations along the long cell axis as a function of cell length in WT (*L3*-*R3* 57509 cells, *ori1* 42612 cells), *ΔmukB* (*L3*-*R3* 27984 cells, *ori1* 54820 cells) and *ΔmatP* (*L3*-*R3* 46679 cells, *ori1* 51350 cells). Sample numbers with different cell lengths are normalized. Cells are oriented to place *L3* more towards the negative pole (towards figure bottom) or, in the *ori1* data, *ter3* is oriented more towards the negative pole (see Fig. S2). White lines denote cell borders. (**E**) Overlay of *ori1* and *L3-R3* localization data in WT and *ΔmatP* from (C) and (D). (**F**) Distance between *L3* and *R3* markers in WT (47376 cells), *ΔmukB* (15615 cells) and *ΔmatP* (41625 cells) in single *L3* and *R3* focus cells. Mean and dispersion (SD) between cells are shown for each distribution. * and ** denote two-sample t-test of *L3*-*R3* distances between WT and *ΔmukB* (p-value 0.0081) and WT and *ΔmatP* (p-value 5 × 10^−4^), respectively. (**G**) (left) Percentage of cells with *L3* and *R3* in opposite cell halves in single *L3* and *R3* focus cells (WT 47376 cells, *ΔmukB* 15615 cells, *ΔmatP* 41625 cells). (right) Percentage of cells with *L3-R3-L3-R3* (or *R3-L3-R3-L3*) configuration (versus *L3-R3-R3-L3* or *R3-L3-L3-R3*) in double *L3* and *R3* focus cells (WT 10352 cells, *ΔmukB* 2535 cells, *ΔmatP* 6297 cells). Scale bars: 1 μm. (**H**) Percentage of cells retaining *L3-R3-L3-R3* orientation (versus flipping to *R3-L3-R3-L3*) from a mother cell to a daughter cell in WT (859 pairs) and *ΔmatP* (1054 pairs). Scale bars: 1 μm. Data from 3 repeats of each experiment.

New-born WT cells exhibited the distinctive *left-oriC-right* (*L3-R3*) chromosome organization (Fig. 2C, D and E, Fig. S2), where *oriC* remained at the cell center and the chromosome arms (*L3* and *R3*) resided in opposite cell halves (97.8 ± 0.6% (±SD), Fig. 2F and G) (Nielsen et al., 2006; Wang et al., 2006). During replication-segregation, the pattern was extended into a translationally symmetric *left-oriC-right-left-oriC-right* (*L3-R3-L3-R3*) pattern in 73.1 ± 3.9% (±SD) of WT cells (Fig. 2G). In the absence of MukBEF, we expected chromosome organization to be more ‘relaxed’ as lengthwise chromosome compaction is relieved (Mäkelä and Sherratt, 2020). Indeed, the localization of all four chromosomal markers was less precise, with a wide distribution of *L3-R3* distances (Fig. 2F), fewer *L3* and *R3* foci localized in opposite cell halves (56.6 ± 3.2% (±SD), Fig. 2G), and a random chance of observing the *L3-R3-L3-R3* organization (47.7± 0.2% (±SD)), versus *L3-R3-R3-L3* or *R3-L3-L3-R3*. The impaired chromosome organization in *ΔmukB* cells is frequently accompanied by the chromosome arms being aligned together along the long cell axis with *ori1* towards the old cell pole (Fig. 2B and D)(Danilova et al., 2007), an organization reminiscent of the situation in wild-type *C. crescentus* (Wang et al., 2013). We conclude that absence of lengthwise compaction by MukBEF axial cores causes loss of both the distinctive *L-R* chromosome organization prior to replication and the *L-R-L-R* organization after replication.

Absence of MatP leads to the formation of circular MukBEF axial cores, rather than linear ones (Fig. 2A). We observed that Δ*matP* cells exhibited chromosome locus localization patterns strikingly different from that of WT and *ΔmukB* cells (Fig. 2 C and E). The average distance between *L3* and *R3* was reduced two-fold (Fig. 2F), consistent with MukBEF-mediated lengthwise compaction of *ter* in the absence of MatP (Mäkelä and Sherratt, 2020). Lengthwise compaction of *ter* reduced the efficiency of *L3* and *R3* being directed into opposite cell halves (65.7 ± 0.8% (±SD), Fig. 2G). Concomitantly, it also led to *L3* and *R3* foci being preferentially localized closer to the cell center than in WT cells where *L3* and *R3* localize towards the cell poles (Fig. 2E). Surprisingly, despite these substantial perturbations, the normal *L3-R3-L3-R3* organization was retained in Δ*matP* cells prior to cell division (80.2 ± 1.9% (±SD)).

Because the circular MukBEF axial cores of Δ*matP* cells lead to rotational chromosome symmetry (Mäkelä and Sherratt, 2020), we hypothesized that non-replicating chromosomes in new born cells are free to rotate around the longitudinal axis of the cell. This would switch the configuration from *L3-R3* to *R3-L3* (or *vice versa*); while bi-lobed replication intermediates could prevent the rotation as replication progresses. To test this hypothesis, we followed Δ*matP* cells under the microscope to observe how often the *L3-R3-L3-R3* orientation flips to *R3-L3-R3-L3* (or *vice versa*) in consecutive generations (Fig. S2). Indeed, *ΔmatP* cells retained the chromosome orientation only in 32.2 ± 4.6% (±SD) of daughter cells, while WT cells predominantly retained the orientation (91.4 ± 5.2% (±SD), Fig. 2H). The *ΔmatP* daughter cells with flipped chromosome orientation were initially born with the same orientation as in the mother cell (88.4 ± 2.8% (±SD), Fig. S2E), indicating that the chromosome rotation generally occurs after division but prior to duplication of the chromosome. We propose that the *L3-R3-L3-R3* organization arises from an interplay between bidirectional replication and the action of MukBEF axial cores.

### DnaN marks lagging strand segregation to the cell center

Translational symmetry of sister chromosomes arises at least in part during DNA replication from the symmetric segregation of lagging strands towards midcell (and leading strands towards the cell poles), as shown using an elegant genetic system (White et al., 2008). Here, we sought directly to visualise the positioning of lagging strands in WT, *ΔmatP* and *ΔmukB* cells.

During replication, ∼40 DNA-bound β_2_-clamps, which ensure DNA polymerase III processivity, have a ∼3 min residence time on DNA before they are unloaded (Moolman et al., 2014). The DNA-bound clamps are expected to accumulate largely on the lagging strand and its template because new clamps are loaded during synthesis of each Okazaki fragment (Fig. 3A). We reasoned that since β_2_-clamps could potentially cover >100kb of newly replicated lagging strand DNA, they could serve as a marker to monitor lagging strand segregation (Fig. 3B). As a reference, for the localization of replication forks, we imaged fluorescent DNA polymerase III ε-subunits (DnaQ). Indeed, while DnaQ foci were more spread towards cell poles as previously described (Reyes-Lamothe et al., 2008), DnaN foci localized closer to the cell center cell center, consistent with the lagging strands being directed to the cell center (Fig. 3C). By directly measuring the distance from each DnaQ focus to the closest DnaN focus, we found that 41.2 ± 5% (±SD) of DnaQ foci do not colocalize with DnaN foci during replication (Fig. 3D). Differential location of bulk DnaN and replication forks was confirmed by measurement of the distances from replicative helicase (DnaB) foci to their closest DnaN focus (47.1 ± 6.1% (±SD) not colocalizing) (Fig. S3B, C). Since DnaN and DnaQ colocalize during early and late replication, when sister replisomes are necessarily close together, we also analysed the localization patterns in mid-replication cycle (Fig. 3E), when independently tracking replication forks are more frequently spatially separate. The pattern of DnaQ foci that did not colocalize with DnaN foci (Fig. S3F) underlines the conclusion that spatially separate sister replisomes in opposite cell halves have a different cellular location from DnaN. Our results are consistent with previous independent measurements of DnaQ and DnaN localization, and the observation that DnaN foci of sister replisomes often do not spatially separate (Mangiameli et al., 2017; Reyes-Lamothe et al., 2008; Wallden et al., 2016). Here we provide the first direct evidence that the replisome and β_2_-clamps frequently do not colocalize during replication.

**Fig. 3.**
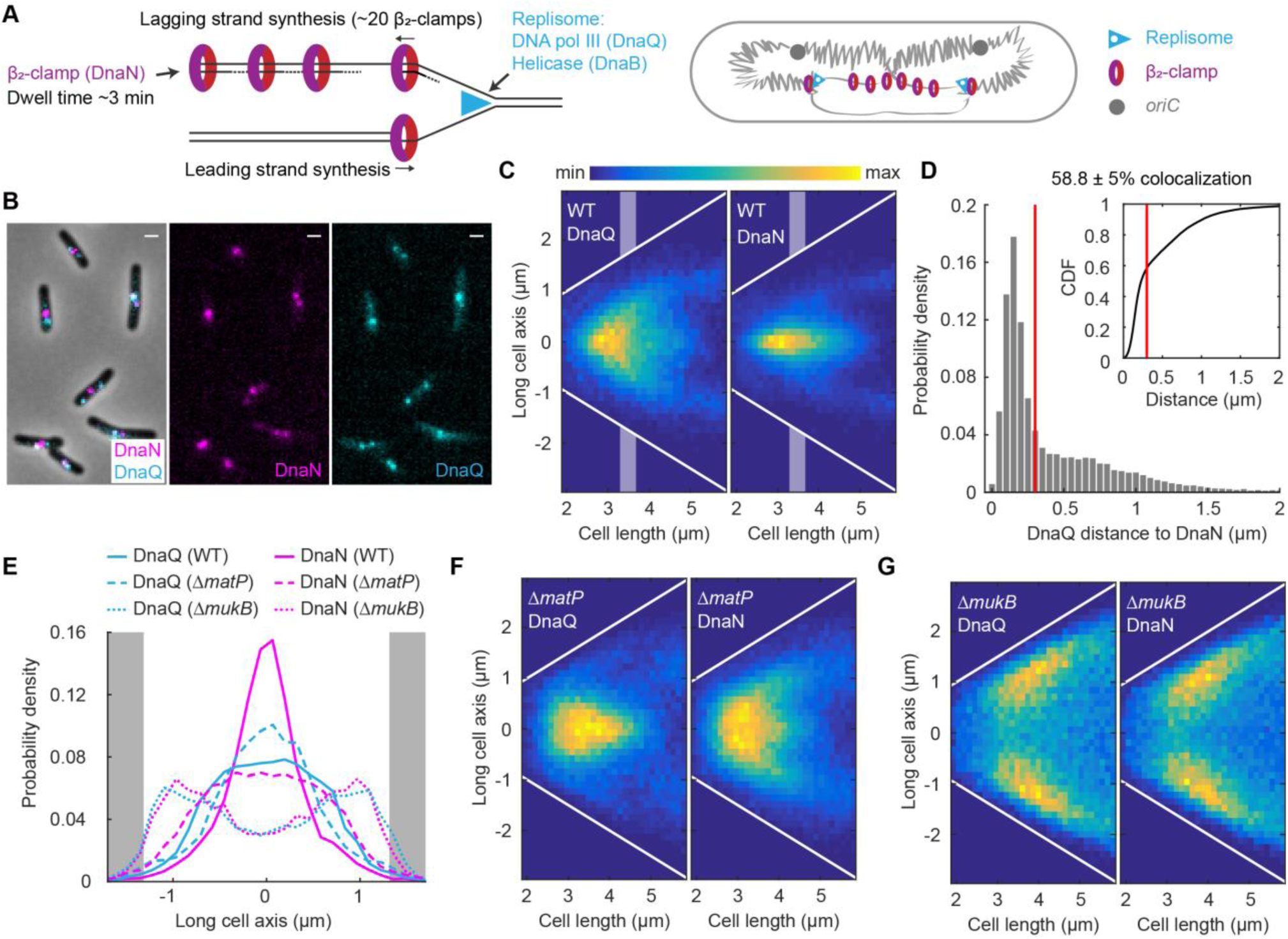
DnaN visualizes the lagging strands during replication. (**A**) Schematic of accumulation of β_2_-clamps (DnaN) on the lagging strand and its template during replication (Moolman et al., 2014). The DNA polymerase ε-subunit (DnaQ) marks the location of the replisome. (**B**) Representative images of WT cells with fluorescently labeled DnaN and DnaQ. Scale bars: 1 μm. (**C**) DnaQ and DnaN localization in WT cells as a function of cell length (37720 cells). White lines denote cell borders. Shaded areas denote intermediate cell lengths for localization data in (E). (**D**) Distance from a DnaQ focus to the closest DnaN focus. DnaQ and DnaN colocalize in 58.8 ± 5% (±SD) of focus pairs (38855 pairs) as defined by a threshold (red lines) below which two proteins colocalize (dictated by a diffraction limit of 300 nm). Inset shows the same data as a cumulative distribution. Same data as in (C). (**E**) DnaQ or DnaN localization with intermediate cell lengths (3.3-3.7 μm) in WT (DnaN 7104, DnaQ 8006 spots), *ΔmatP* (DnaN 11925, DnaQ 8025 spots) and *ΔmukB* (DnaN 5060, DnaQ 4205 spots) cells (see C, F, G). Full width at half maximum (FWHM) (±SD) of the distribution in WT: DnaN 0.67 ± 0.06 μm, DnaQ 1.67 ± 0.08 μm; and in *ΔmatP*: DnaN 1.85 ± 0.04 μm, DnaQ 1.14 ± 0.14 μm. Grey areas denote cell poles. (**F**) DnaQ and DnaN localization in *ΔmatP* cells (51956 cells) and in (**G**) *ΔmukB* cells (22902 cells) as a function of cell length. White lines denote cell borders. Data from 3 repeats in all analyses.

Our direct visualization of the segregation of lagging strands during replication, supports the previously shown symmetric segregation of leading strands towards the cell poles (White et al., 2008). To analyze how MukBEF and MatP contribute to lagging strand segregation, we measured DnaN localization in *ΔmatP* and *ΔmukB* cells. The DnaN distribution in *ΔmatP* cells was much broader than in WT cells (Fig. 3E and F), indicative of spatially less precise lagging strand segregation, but still directed towards cell centers, as predicted by the *L3-R3-L3-R3* organization. The DnaQ distribution in mid-cycle *ΔmatP* cells was more central than that of DnaN (50.8 ± 1.3% (±SD) colocalization with DnaN, Fig. 3E), most likely because of less separated chromosome arms. Both DnaQ and DnaN exhibited a broader distribution at shorter cell lengths (Fig. 3F), presumably because of a more random chromosome conformation (Fig. 2). *ΔmukB* cells showed a bimodal distribution of DnaN and DnaQ localizations towards cell poles, with almost identical patterns for both markers (Fig. 3G). This shows that lagging strands and their templates cannot be directed to cell centers in a timely manner in the absence of MukBEF function, a result consistent with impaired *L-R* and *L-R-L-R* organization in Δ*mukB* cells. By measuring the distance from each DnaQ focus to the closest DnaN focus, we found that lagging strands did not leave the vicinity of the replisome during the DnaN dwell time on chromosomes of ∼3 min (78.4 ± 0.5% (±SD) of foci; Fig. S3G). We hypothesize that this is a consequence of delayed decatenation by TopoIV in the absence of MukBEF (Zawadzki et al., 2015), since lagging strand templates can only be segregated from the leading strands once decatenation has occurred.

Finally, a dynamin-like protein CrfC (aka YjdA) has been proposed to bind β_2_-clamps and tether the nascent strands of sister-chromosomes together (Ozaki et al., 2013). However, upon deletion of *crfC*, we observed no changes to DnaN localization along the long cell axis, or any decrease in the frequency of the *L3-R3-L3-R3* configuration (Fig. S3G, H). This result indicates that CrfC is not necessary for WT chromosome conformation and segregation.

### Ancestral DNA strands are preferentially retained at older cell poles

Previously it has been hypothesized that a symmetrical segregation of lagging strands to the cell center leads to sister chromosomes’ translational symmetry and, in consequence, the ancestral (‘immortal’) template DNA strand is not randomly segregated to daughter cells over subsequent generations but preferentially retained in the daughter with the older cell pole (discussed in (Toro and Shapiro, 2010)). Cell division generates two new cell poles at the division septum, while the other ends of the daughter cells are the older poles that were created in an earlier division. To address this theory directly, we have developed a novel pulse-chase assay. It allowed us to visualize relative age of DNA strands between sister chromosomes and relate their position to the age of the pole without the need for cell synchronization or tracking (Fig. 4A).

**Fig. 4.**
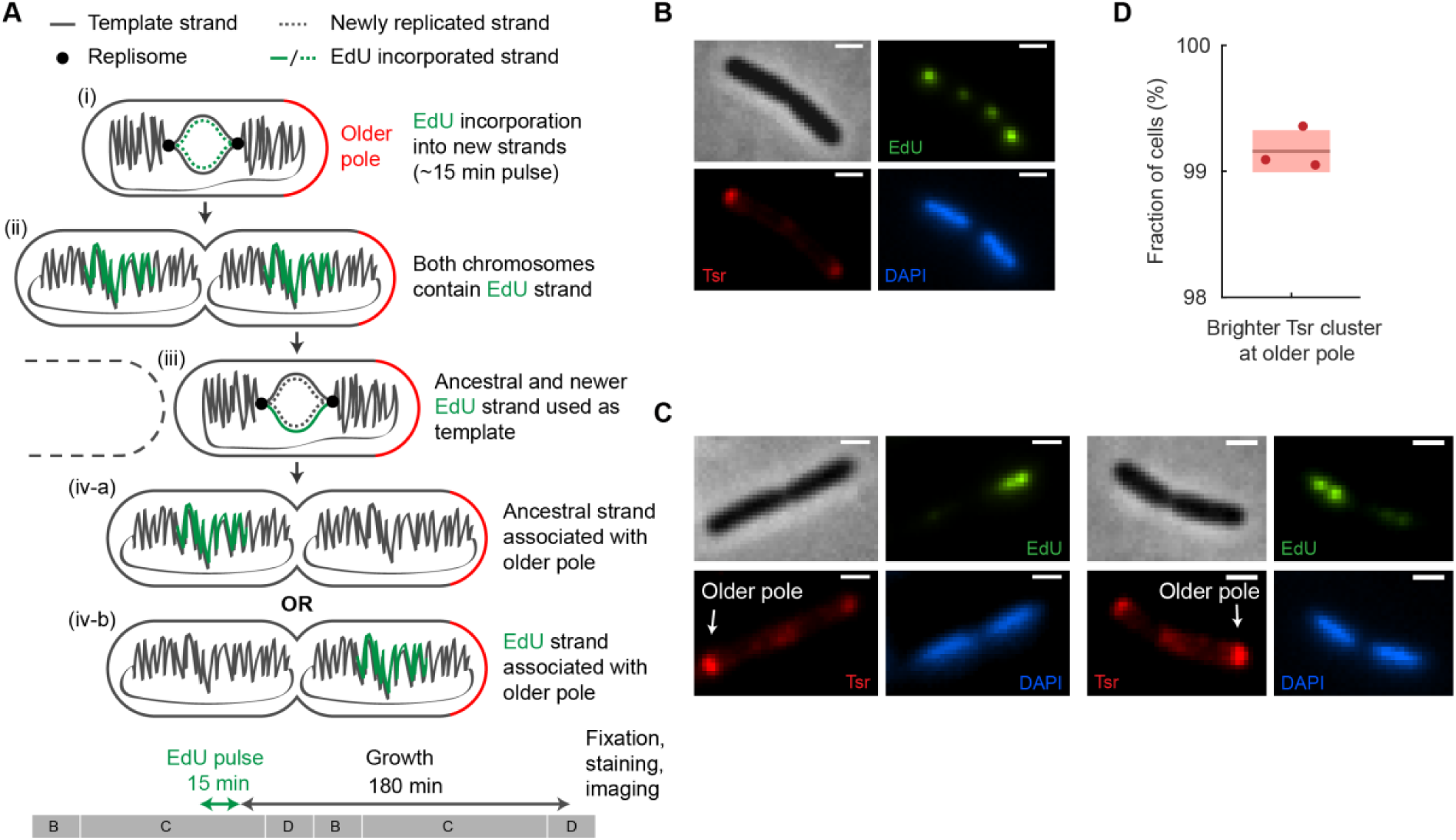
Visualization of ancestral DNA strand retention in *E. coli*. (**A**) Ancestral DNA strand propagation shown following an EdU pulse and the subsequent growth. After the 2^nd^ round of replication only one of the chromosomes inherits the EdU label. Note that only a part of the chromosome is labelled with EdU. The 15 min EdU pulse and growth period are also shown relative to a schematic of cell cycle stages (B, C and D periods, generation time ∼150 min). (**B**) Representative EdU^Alexa488^, Tsr^TMR^ and DAPI images of a cell at stage (ii) (see (A)) after the EdU pulse. Note that each chromosome has two EdU foci because pulse-labeled chromosome arms are separated. Scale bars: 1 μm (**C**) Representative EdU^Alexa488^, Tsr^TMR^ and DAPI images after the complete pulse-chase protocol (stage (iv-a), see (A)). The older pole is indicated with an arrow. Scale bars: 1 μm. (**D**) Accuracy of the older pole classification using Tsr prior to cell division. Shaded areas denote SD. Data from 2505 cells and 3 repeats.

The assay comprises pulse labelling of newly replicated DNA and identifying relative pole age by chemoreceptor accumulation at cell poles. The newly synthesized DNA was labelled by a 15 min EdU (5-Ethynyl-2’-deoxyuridine) pulse, after which cells were washed, and allowed to grow for 3 h (generation time ∼150 min). To avoid EdU-mediated growth defects, thymidine was added to the medium to outcompete EdU. We observed no detrimental effects on growth rate or cell size from the low concentration of EdU used in the pulse (Fig. S4). After the growth period, which was longer than a single generation time, most cells have completed an additional round of replication resulting in only one of the two sister chromosomes remaining EdU-labelled (Fig. 4A). Cells were fixed and EdU was visualized by click-chemistry using Alexa 488 azide. In cells with segregated chromosomes just before division, the chromosome with new strand was fluorescently labeled, while the one with the ancestral strand was not.

To identify the older cell pole, we exploited the fact that the serine chemoreceptor, Tsr, accumulates approximately linearly with time at the cell poles (Ping et al., 2008). Hence the older pole can be distinguished from the new pole by a higher quantity of fluorescently labeled Tsr. Because imaging the Tsr-GFP fusion used before (Ping et al. 2008) was incompatible with EdU staining, we devised an alternative labeling method. To this end, a functional HaloTag fusion of the endogenous *tsr* gene was labeled with synthetic TMR dye (Fig. 4B). Furthermore, nucleoids were stained by DAPI. This allowed us to determine if the older strand chromosome was segregated towards the older or newer pole in each cell (Fig. 4C). We limited analysis only to cells with segregated chromosomes (i.e. two visibly separate nucleoid regions) only one of which displayed EdU fluorescence. Note that this does not bias the analysis towards a specific pole. In a control experiment we confirmed that the intensity of Tsr-mYpet foci was higher at the older pole in 99.2 ± 0.5% (±SD) of cells.

We observed that 71.3 ± 3.9% (±SD) of WT cells contained EdU foci in the chromosome closer to the new pole (Fig. 5A). Because EdU was incorporated into the new template strand, this indicates that the ancestral strand is preferentially retained at the older pole. The result deviates significantly from random retention, where the older pole would have a 50% chance of inheriting either strand (binomial two-tailed test p-value < 10^−5^). We also compared the dispersion (SD) of our data to a binomial distribution with different sample sizes to estimate reliability of our experiment (Fig. 5B). We found excellent agreement showing that our measurements are robust for the given sample size, with no additional noise sources, and increasing data sample size would give diminishing returns.

**Fig. 5.**
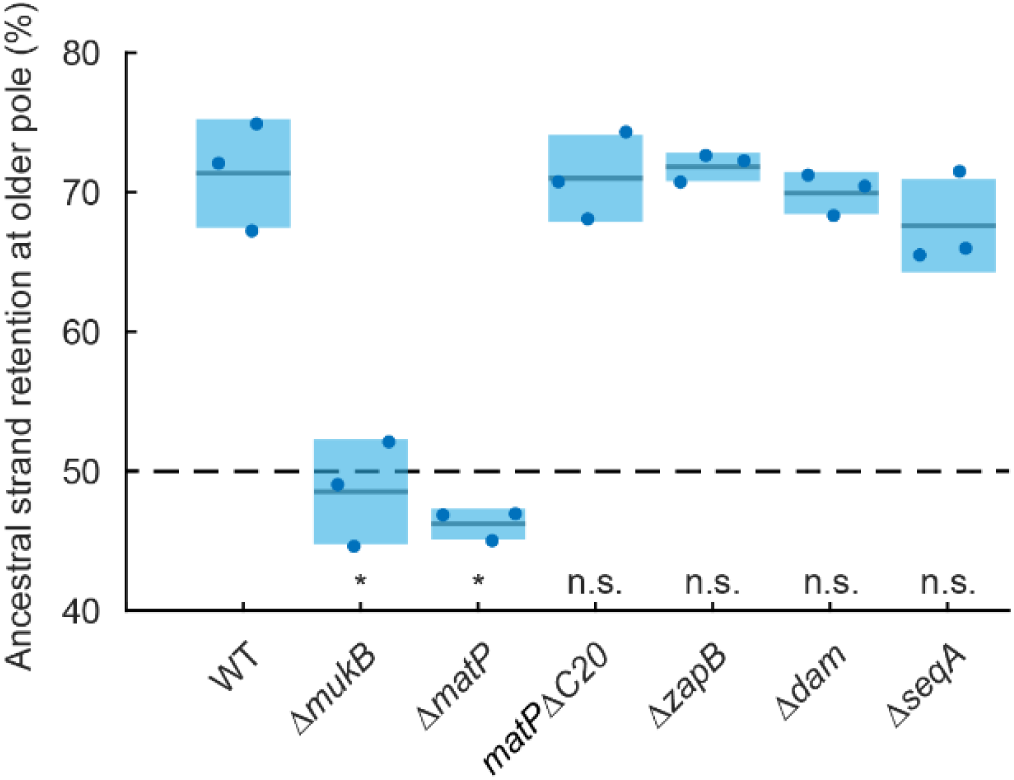
Preferential retention of the ancestral strand at older cell poles requires functional MukBEF and MatP. Percentage of ancestral strands retained at the older pole in WT (988 cells), *ΔmukB* (427 cells), *ΔmatP* (1050 cells), non-divisome interacting *matPΔC20* mutant (1617 cells), *ΔzapB* (969 cells), *Δdam* (717 cells) and *ΔseqA* (473 cells). Dashed line shows random retention. p-value from two-proportion two-tailed z-test was used to test if binomial distributions significantly differ from WT; indicated by n.s. (> 0.01, non-significant) and * (< 0.01, significant) (p-values <10^−5^, <10^−5^, 0.51, 0.83, 0.26, and 0.03, respectively). Shaded areas denote SD. Data from 3 repeats.

How is ancestral strand retention related to chromosome organization? To address this question, we tested the contributions of MukBEF and MatP to ancestral strand retention. Upon deletion of *mukB*, we observed a random segregation of the ancestral strand (48.5 ± 3.8% (±SD), Fig. 5A), demonstrating that functional MukBEF is required for ancestral strand retention at older poles. Deletion of *matP* also abolished the preferential segregation of the ancestral strand (46.2 ± 1.1% (±SD), Fig. 5A). While MatP has not been implicated in early chromosome segregation, when the segregation pattern(s) emerge, the influence of MatP on MukBEF action is crucial as it prevents longitudinal chromosome rotation (Fig. 2H), which would disrupt the association of the ancestral strand with the older pole. MatP-*matS* also interacts with the divisome through ZapB and this interaction has been proposed to partially anchor *ter* to the inner cell membrane (Espéli et al., 2012). This interaction could plausibly contribute to the ancestral strand retention by anchoring the chromosome and thereby preventing chromosome rotation. However, upon replacing the native *matP* with a non-divisome interacting *matPΔC20* mutant or deleting *zapB*, we did not observe any difference to WT with regard to ancestral strand retention (71.0 ± 3.1% and 71.8 ± 1.0%, respectively (±SD); Fig. 5A). This confirms that the loss of ancestral strand retention in *ΔmatP* cells is related to the proposed longitudinal rotation of the chromosome over generations.

Finally, since MukBEF and MatP have co-evolved with a group of proteins (including Dam and SeqA) that are related to Dam DNA methyltransferase activity (Brézellec et al., 2006), we tested the influence of these proteins on the retention of the ancestral strand. Dam methylates adenines in the sequence GATC, which transiently distinguishes strands after replication because of delayed methylation of the newly replicated strands. Prior to Dam methylation, SeqA binds to hemimethylated GATC sites, negatively regulating replication initiation and possibly contributing to chromosome segregation (reviewed in (Waldminghaus and Skarstad, 2009)). Deletion of either *dam* or *seqA* did not influence ancestral strand retention at older poles (69.9 ± 1.5% and 67.6 ± 3.3%, respectively (±SD); Fig. 5A), indicating GATC methylation patterns do not affect the observed asymmetry and consequently, overall *L*-*R* chromosome organization.

## Discussion

Our results demonstrate how MukBEF axial cores direct nucleoid organization and non-random segregation of sister chromosomes in *E. coli*. This work extends that describing chromosome organization by MukBEF axial cores (Mäkelä and Sherratt, 2020), and uncovers important principles of nucleoid organization and replication-segregation, as illustrated in Fig. 6 and outlined below.

1. MukBEF axial cores, linearized by *matS*-MatP-mediated depletion of MukBEF from *ter*, are required for *left-oriC-right* chromosome locus organization along the long cell axis.
2. The linearity of axial cores prevents longitudinal nucleoid rotation, thereby retaining the *left-oriC-right* chromosome orientation over generations.
3. Translational symmetry of the sister chromosomes results from the axial core structures together with the segregation of lagging strands towards the cell center.
4. The ancestral DNA strand is preferentially segregated to the older pole cell over generations, a process dependent on MukBEF and MatP.
5. Absence of MukBEF leads to anucleate cell formation predominantly at the newer mother cell pole, largely because of failure to segregate newly replicated origins from the vicinity of the older pole.

**Fig. 6.**
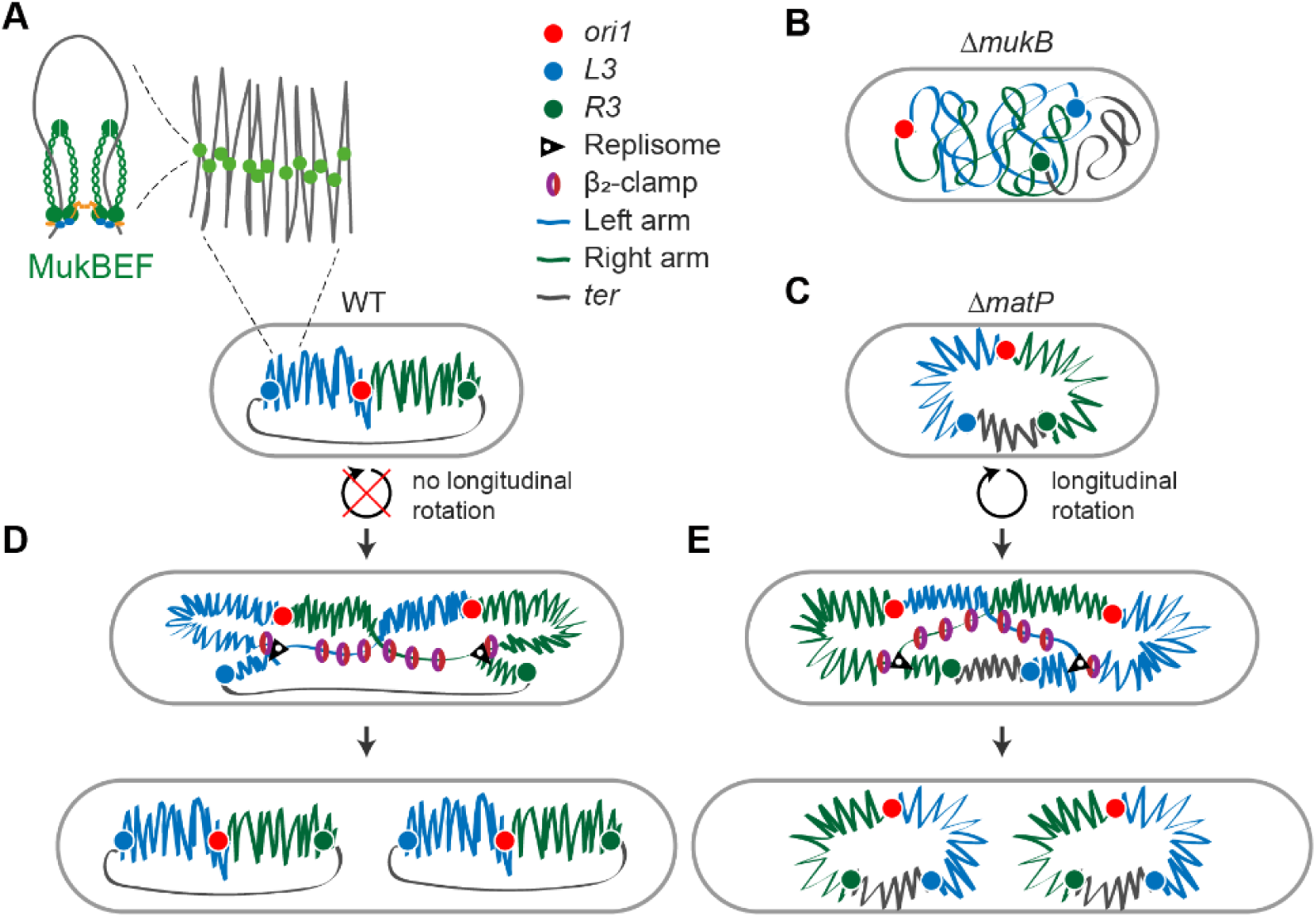
Chromosome organization and segregation by MukBEF and MatP. (**A**) MukBEF dimer of dimer complexes form DNA loops dependent on ATP hydrolysis. Ubiquitous loop formation outside of *ter* compacts the chromosome lengthwise forming a linear axial core to the chromosome. Axial cores form stiff linear nucleoid bundles that localize chromosomal loci linearly along the long cell axis and maintain chromosome orientation by preventing longitudinal cell axis rotation. (**B**) Absence of functional MukBEF increases the effective contour length of chromosomes, leading to mis-localization of chromosome loci and loss of *left-oriC-right* organization of the chromosome. (**C**) MukBEF forms a complete circular axial core to the chromosome in the absence of MatP. Consequently, chromosome arms cannot be efficiently directed to opposite cell halves, *left-oriC-right* organization in non-replicating cells is impaired and chromosomes rotate in the longitudinal cell axis. During replication and prior to division, WT (**D**) and *ΔmatP* cells (**E**) exhibit translational symmetric (*L-R-L-R*) segregation of sister chromosomes. This is accompanied by the symmetric segregation of lagging strands and their templates during replication, as visualized by accumulation of β_2_-clamps. Relative FROS marker, replisome and β_2_-clamps localizations are derived from the data generated here.

Our analyses provide molecular mechanisms underlying the *E. coli* chromosome organization and segregation, and complement previous studies that have rigorously quantified the nucleoid dynamics in mechanical terms (Cass et al., 2016; Fisher et al., 2013; Fritsche et al., 2012). The model succinctly explains previous observations and provide a conceptual foundation for understanding how nucleoid conformation and dynamics shape the subcellular organization.

### E. coli chromosome organization

Stiff nucleoid ‘bundles’ that are radially confined by cell dimensions and exhibit a contour length of the scale of cell dimensions were characterized in live-imaging studies of WT *E. coli* (Fisher et al., 2013). Bundles were also identified in cells with increased volume, which allowed visualisation of non-replicating toroidal chromosomes (Wu et al., 2019). We propose that MukBEF axial cores correspond to these bundles; the duality of the bundles observed in WT cells likely relate to the bimodal nature of axial cores in non-replicating MatP^+^ cells. We hypothesize that individual dimer of dimer MukBEF complexes within axial cores form DNA loops using energy from ATP, most likely through progressive loop enlargement (Davidson et al., 2019; Ganji et al., 2018; Goloborodko et al., 2016). Unloading and re-loading of MukBEF complexes will impact nucleoid spatio-temporal dynamics and could be dependent on intra-chromosomal stress (Fisher et al., 2013). The correlative timescale of bundle dynamics is of the order of a minute, similar to the dwell time of individual MukBEF complexes on the chromosome (Badrinarayanan et al., 2012; Fisher et al., 2013; Mäkelä and Sherratt, 2020). MukBEF forms a linear axial core to the chromosome by compacting it lengthwise outside of *ter*, where *matS* bound MatP displaces MukBEF (Mäkelä and Sherratt, 2020). The absence of MatP leads to formation of circular symmetric MukBEF axial cores. Theoretical studies concluded that lengthwise compaction of the chromosome (by MukBEF) would maintain the linear organization and forms rigid chromosome bundles (Marko and Siggia, 1997). We propose that the linearity of the axial core reduces the dimensions of free chromosome movement, thereby preventing longitudinal chromosome rotation and therefore explaining how chromosome loci can be robustly positioned along the long cell axis in a colinear manner outside of *ter* (Fig. 6).

Previous attempts to explain chromosome locus positioning by a randomly oriented polymer, or by transcription factor-mediated DNA loops fail to explain nucleoid organization in Δ*mukB* or Δ*matP* cells (Fritsche et al., 2012; Jun and Mulder, 2006). The presence of MukBEF axial cores explains how loss of a flexible *ter* region in the absence of MatP results in chromosome arms being closer together, as shown here. A flexible *ter* region might be required for efficient chromosome segregation during fast growth as Δ*matP* cells exhibit more frequent anucleate cell production than MatP^+^ cells (Mercier et al., 2008). Broadly, we propose that the rigid linear axial core removes the requirement for membrane anchoring to orient and/or position the chromosome. Membrane tethering is generally found in organisms in which MukBEF has been replaced by Smc-ScpAB complexes and which carry a *parABS* segregation system; e.g. through PopZ in *C. crescentus* (Ebersbach et al., 2008), HubP in *V. cholera* (Yamaichi et al., 2012), and RacA/DivIVA in sporulating *B. subtilis* (Ben-Yehuda et al., 2003; Wu and Errington, 2003). Membrane anchoring typically uses ParB bound to *oriC*-proximal *parS* sites as an intermediary. Intriguingly, some bacteria, such as *V. cholera* or *P. aeruginosa*, not only encode MukBEF/MksBEF, but also specify a *parABS* system (David et al., 2014; Vallet-Gely and Boccard, 2013). Whether organisms that encode MukBEF orthologs, but not typical Smc-ScpAB complexes, and which lack ParABS systems, generally have life cycles that encompass overlapping replication cycles, similar to *E. coli*, remains to be determined.

### Sister chromosome replication and segregation

Chromosome organization specified by the MukBEF axial core directs the translational symmetric (*L-R-L-R*) segregation of sister chromosomes during replication (Fig. 6), which alone is insufficient for this organization (Fig. 6). Lengthwise compaction of newly replicated DNA promotes individualization of sister chromosomes through excluded volume interactions and by maximization of conformational entropy that leads to repulsion between sister chromosomes (Goloborodko et al., 2016; Marko and Siggia, 1997). The presence of the *L-R-L-R* segregation pattern of sister chromosomes in both WT and Δ*matP* cells indicates that the pre-replication organization of the chromosome, where chromosome arms are directed into separate cell halves only in WT cells, is not required for establishing this pattern. While symmetrically lengthwise compacted chromosomes in Δ*matP* cells (Fig. 2A)(Mäkelä and Sherratt, 2020) are prone to change *L-R* orientation in non-replicating cells, bi-loped replication intermediates prevent rotation during late replication (Fig. 6).

The relationship between *L-R-L-R* chromosome organization and symmetrical segregation of leading/lagging strands has been discussed previously in relation to the observed leading strand spatial segregation pattern (Toro and Shapiro, 2010; White et al., 2008). Consistent with this, we observed accumulation of β_2_-clamps, present primarily on lagging strands, towards cell centers of replicating cells, when compared to both DNA polymerase III and helicase localization. Differential positioning of the replisome and β_2_ clamps resolves the conundrum that emerged from studies that favoured a model of a single replication ‘factory’ containing two replisomes at cell center, based on clamp labelling (Mangiameli et al., 2017). The results support the model of independent tracking of the two often spatially separated replisomes in cells undergoing a single round of replication (Japaridze et al., 2020; Reyes-Lamothe et al., 2008) although segregation forces along with the reorganisation of parental and newly replicated DNA leads to frequent movement of sister replisomes towards cell center. A dynamin-like protein YjdA (aka CrfC), a possible candidate for directing β_2_-clamps to the cell centre, was not required for this action. Because clamp localization at the cell center is dependent on the formation of linear axial cores, we hypothesize that MukBEF could plausibly differentiate the strands, e.g. leaving lagging strands less compacted (Fig. 6). However, the different contributions of MukBEF activity to chromosome organization and segregation are difficult to delineate, as in the absence of MukBEF, cohesion time is increased and *L-R* organization prior to replication is impaired (Fig. 1).

Our results also show that the lifetime of individual chromosome-associated clamps (estimated to be ∼3 min (Moolman et al., 2014)) must be longer than the daughter chromosome cohesion time for chromosomal regions outside of *oriC* and *ter* (estimated ∼14 min and ∼9 min, respectively) (Nolivos et al., 2016; Wang et al., 2008). Cohesion time is largely determined by the time required for TopoIV to remove replicative catenanes (Nolivos et al., 2016; Wang et al., 2008; Zawadzki et al., 2015). In addition, slower *oriC* segregation may additionally require accumulation of sufficient newly replicated DNA that leads to abrupt separation of newly replicated sister chromosomes (Cass et al., 2016; Fisher et al., 2013). Cohesion time is influenced by the absence of MukBEF, which promotes TopoIV activity (Nolivos et al., 2016; Zawadzki et al., 2015), although tethering of *ter* to the divisome through MatP-ZapB interactions may also influence cohesion time in this region (Monterroso et al., 2019). Precise measurements of cohesion times for the rest of the chromosome have been refractory to precise experimental determination.

### Ancestral strand retention at the older pole

We have directly shown preferential retention of the ancestral DNA strand at the older pole in *E. coli*. This non-random segregation is determined by the translational symmetry of the sister chromosomes (*L-R-L-R*), along with efficient maintenance of chromosome orientation over generations. Intriguingly, in the absence of MukBEF, preferential strand retention is lost and chromosomes adopt a longitudinal *oriC*-*ter* chromosome organization with co-aligned chromosome arms along the long cell axis, similar to *C. crescentus* and sporulating *B. subtilis*. However, as *E. coli* lacks the properties of cell differentiation, development and regeneration of a multicellular organism, it is not clear why it has evolved a chromosome organization that preferentially segregates the ancestral DNA strand to the cell with the older pole. *E. coli* older pole containing cells exhibit a constant growth rate for hundreds of generations (Wang et al., 2010). However, the death rate was found to increase with replicative cell age, which was attributed to growth-independent accumulation of protein damage (Wang et al., 2010). Increasing cellular maintenance processes through the general stress response reduced the death rate while its absence increased it (Yang et al., 2019). The older pole accumulates more membrane proteins (e.g. chemoreceptors, efflux pumps) than the new pole and in fluctuating or poor environment, these can significantly contribute to cell growth (Bergmiller et al., 2017; Ping et al., 2008). For example, the main multidrug efflux pump of *E. coli*, AcrAB-TolC, exhibits a partitioning bias for the older cell poles (Bergmiller et al., 2017). Consequently, older pole cells display increased efflux activity relative to new cell pole daughters giving the older pole cell a growth advantage under subinhibitory antibiotic concentrations and possibly protection against other toxic compounds. Notably, AcrAB-TolC pump activity is also required for acquiring a resistance gene from mobile genetic elements in the presence of antibiotics, as it reduces antibiotic concentrations inside the cell (Nolivos et al., 2019). A common epigenetic mechanism to regulate phase variation in bacteria involves formation of DNA methylation patterns by proteins binding near a hemimethylated GATC site, and blocking methylation, e.g. *pap* or *foo, clp*, and *pef* systems, which all encode pili (Casadesús and Low, 2013). Preferential retention of the old strand at the old pole could potentially cause the old pole cell to more likely to maintain the previous methylated state. We also hypothesize that ancestral strand retention at older pole cells could be beneficial to structured growth of *E. coli* in colonies where younger bacteria could progress to a new terrain while the older ones stay closer to the colony center. Finally, older strand retention could simply be an evolutionary by-product of maintaining the *left-oriC-right* chromosome organization over division cycles. Nevertheless, since ancestral strand retention occurs in only ∼70% of older-pole cells, this gives opportunities for selection in fluctuating or harmful environments independent of whether older or newer pole cells thrive better.

## Materials and Methods

### Bacterial strains and growth conditions

Bacterial strains and primers are listed in Table S1 and S2, respectively. All strains were derivatives of *E. coli* K12 AB1157 (Bachmann, 1996). *kan, cat, gen*, and *hyg* refer to insertions conferring resistance to kanamycin (Km^r^), chloramphenicol (Cm^r^), gentamycin (Gm^r^) and hygromycin B (Hyg^r^), respectively. The insertions are flanked by Flp site-specific recombination sites (*frt*) that allow removing the resistance gene using Flp recombinase from plasmid pCP20 (Datsenko and Wanner, 2000). *tsr-HaloTag-kan* and *tsr-mYpet-kan* were inserted into the native *tsr* chromosomal locus using λ-red recombination (Datsenko and Wanner, 2000). The generated gene loci were transferred by phage P1 transduction to AB1157 yielding strains JM122 (*tsr-HaloTag-kan*) and JM133 (*tsr-mYpet-kan*). Deletion strains of JM122 were constructed by P1 transduction, first removing the *kan* resistance gene using Flp recombinase. *L3-R3* deletion strains were constructed from RRL66 using P1 transduction. The microfluidics strain (JM09) was constructed from RRL189 by introducing *Δflhd-kan* and *ΔmukB-kan* by consecutive rounds of P1 transduction and Flp recombination. GFPmut2 cell marker was inserted at an *attTn7* site by a plasmid transformation as described in (McKenzie and Craig, 2006). The DnaQ and DnaN labeled strain (JM141) was constructed from RRL388 using P1 transduction from RRL36. Deletion strains of JM141 were constructed by P1 transduction, first removing the *kan* resistance gene using Flp recombinase. JM142 and JM143 were constructed by P1 transduction from JW4070. All genetic modifications were verified by PCR and/or sequencing and behavior in quantitative imaging. *mukB* deletions were verified by temperature-sensitivity in rich media, as described in (Nolivos et al., 2016).

Cells were grown in M9 minimal medium supplemented with 0.2% (v/v) glycerol, 2 μg ml^-1^ thiamine, and required amino acids (threonine, leucine, proline, histidine and arginine; 0.1 mg ml^-1^) at 30 °C. For microscopy, cells were grown overnight, diluted 1000-fold and grown to an A_600_ of ∼0.1. Cells were then pelleted, spotted onto an M9 glycerol 1% (w/v) agarose pad on a slide and covered by a coverslip. In mother machine microfluidics experiments, cells were first grown in M9 minimal medium with 0.2% (v/v) glycerol at 30 °C (as above), and then placed inside the microfluidics device, when the media was changed to M9 minimal medium supplemented with 0.2% (v/v) glucose, 2 μg ml^-1^ thiamine, MEM amino acids (Gibco, #11130-036), 0.1 mg ml^-1^ proline, and 0.85 mg ml^-1^ Pluronic F127 (Sigma-Aldrich, P2443), and the temperature was set to 37 °C.

### EdU pulse labeling

Cells grown until A_600_ of ∼0.1 were labelled with 10 μM EdU (5-Ethynyl-2’-deoxyuridine, Thermofisher, C10337) for 15 min after which cells were washed, introduced to fresh media containing 60 μg/ml thymidine and allowed to grow for 3 h. Following this, cells were fixed with 4% PFA (v/v) for 30 min and permeabilized with 0.5% Triton X-100 (v/v) for 30 min. EdU click-chemistry reaction was conducted following the instructions (Thermofisher, #C10337) using Alexa 488 azide in a final volume of 50 μl for 30 min at room temperature, followed by washing. Cells were then labelled with TMR HaloTag ligand as in (Banaz et al., 2019). Briefly, cells were incubated with 2 μM TMR ligand for 30 min and washed several times. Finally, nucleoids were labelled with 1 μg/ml DAPI for 15 min and washed, after which the cells were ready for imaging.

### Epifluorescence microscopy

Fluorescence images were acquired on an inverted fluorescence microscope (Ti-E, Nikon) equipped with a perfect focus system, a 100× NA 1.4 oil immersion objective, a motorized stage, an sCMOS camera (Orca Flash 4, Hamamatsu), and a temperature chamber (Okolabs). Exposure times were 300 ms for TMR, Alexa 488 and mCherry, mYpet; 150 ms for mCerulean, and 100 ms for DAPI using an LED excitation source (Lumencor SpectraX). Phase contrast images were collected for cell segmentation. Microscopy data was collected automatically from the sample area. Time-lapse images were collected every 10 min for 3 h with a shorter exposure time of 150 ms, except fluorescence images from the microfluidics device were collected every 5 min.

### Microfluidic devices

The microfluidic single-cell imaging device (“mother machine”) was prepared as in (Uphoff, 2018). The device was designed using Autodesk AutoCAD software. The dimensions of the cell channels were 1.2 μm x 1.2 μm x 20 μm and the media flow channels were 100 μm x 25 μm. The structures were fabricated on a silicon wafer (Kavli Nanolab, Delft University) (Moolman et al., 2013) and a negative polydimethylsiloxane (PDMS) mold was created from the silicon wafer using a 5:1 mixture of monomer and curing agent (Dow Corning Sylgard 184 Kit). After removing air bubbles using vacuum, the chip was cured at 65°C for 1.5 hours. The mould was treated with Trichloro(1H,1H,2H,2H perfluorooctyl)silane (Sigma) in vacuum overnight. The PDMS device was generated from the negative mold using a 10:1 mixture of monomer and curing agent and cured at 65°C for 1.5 hours. Media flow holes were punched through the device with 0.75 mm diameter. Cover slips were cleaned by sonication in acetone for 20 min, washing with dH2O, sonication in isopropanol for 20 min, and dried with nitrogen. The PDMS device was washed with isopropanol and dried with nitrogen. The device and a cover slip were bonded using air plasma (Plasma Etch PE-50) and placed in an oven at 95°C for 30 min. Cells were pipetted into the device and the device was centrifuged at 4000 rpm for 10 min to place cells into the channels. The media supplemented by Pluronic F127 was fed into the device through silicon tubing (Tygon ND 100-80 microbore, VWR) using a motorized infusion pump (New Era Pump Systems). Initially, a high flow rate of 1.5 ml/hr was applied to flush the excess cells and then lowered to 0.5 ml/hr. After this, cells were allowed to grow for ∼2 h before starting the time-lapse imaging.

### Image analysis

Cell based information, including cell outlines, lineages, pole ages, per pixel fluorescence intensities, and fluorescent marker localization, was extracted using SuperSegger (Stylianidou et al., 2016) in MATLAB (MathWorks). SuperSegger uses an image-curvature method to identify foci to avoid the identification of false positive foci due to background intensity from cytoplasmic fluorescence and uses a gaussian fit to find the subpixel resolution location of foci. Focus quality is determined by a combination of intensity and fitting parameters and bad quality foci were filtered out. A threshold value was confirmed by visual inspection and the same threshold was used for all compared data sets. The channels were aligned prior to analysis.

### Mother machine analysis

From the lineage data, cells were classified as ‘normal’ growing cells, anucleate cells, mothers of anucleate cells and sisters of anucleate cells. Cells that disappeared early or didn’t have a tracked lineage were excluded from the analysis. Anucleate cells were considered as cells that didn’t divide, didn’t elongate and lacked an *ori1* marker present, while its sister cell elongated and divided normally, and had *ori1* marker(s). If neither of the sister cells divided normally, cells were excluded from analysis. The older pole of the anucleate cell was traced back ≥2 generations to determine whether the anucleate cell formed on the older or newer pole of the mother. The cell size at birth (Fig. S1B) was determined at the first frame of each cell and the number of *ori1* foci prior to division (Fig. 1C) at the last frame of a cell.

### Fluorescent marker localization

For *ori1, ter3, L3* and *R3* markers, intensity profiles with different cell lengths were normalized, as the expectation is that a cell will have at least a single focus at all times. As cell orientation is random relative to the pole age, cells were oriented to place *L3* more towards the negative pole than *R3* and, in the *ori1* data, *ter3* was oriented more towards the negative pole. To determine flipping frequency of *L3-R3-L3-R3* markers from time-lapse imaging, first mother cells that contained at *L3-R3-L3-R3* or *R3-L3-R3-L3* were identified. Next, their daughter cells with *L3-R3-L3-R3* or *R3-L3-R3-L3* were identified. The angle (Fig. S2) between vectors pointing from the more polar *L3* to the more polar *R3* was calculated between mother and daughter cells. If the angle exceeded 90°, the chromosome orientation was considered flipped. To measure width of a unimodal distribution and to avoid inaccuracy from binning the data, the data was fitted by a kernel distribution in MATLAB and full width at half maximum (FWHM) was calculated from the fitted distribution.

### EdU pulse labelling

A functional HaloTag fusion of the endogenous *tsr* gene was used in the EdU labeling, as click-chemistry reaction conditions are detrimental to conventional fluorescent proteins. To measure EdU association with the older pole, the following criteria were used to select cells from an asynchronous cell population (see Fig. S4). First, Otsu’s thresholding (Otsu, 1979) was used to segment nucleoid area(s) from the cellular background and only cells that have two separate, large-enough nucleoid areas were analysed. Second, a cell must exhibit a clear difference in polar Tsr intensity. The center line of a cell was extracted to find coordinates of cell poles by fitting a cell mask to a second-order curve. The intersection of the cell mask border and the curve was used to define cell poles. Median Tsr intensity of the cell area was subtracted from all Tsr pixel intensities and a sum of 9 brightest pixels from each pole were used to quantify the pole intensity. To minimize effects of noise and discrete pixel size in segmentation, only cells with >1.5-fold difference in polar Tsr intensity were analysed. The pole that had a higher intensity of Tsr was designated as the older pole. Third, only one of the nucleoids must be labelled by EdU. EdU with short incorporation times appear as distinct foci (see Fig. 4 and Fig. S4). The foci below a fixed threshold for the score were discarded. The foci were mapped to the nucleoids by projecting coordinates of both on the center line of the cell. With these criteria, the processed microscopy data from SuperSegger was automatically analysed to extract the result of EdU association with older cell pole. To avoid segmentation errors of the cell area, correct cell segmentation was visually inspected and inaccurately segmented cells were removed.

### Tsr time-lapse

The accuracy of Tsr-based identification of the old cell pole in our growth conditions was estimated by tracking cells with a functional mYpet fusion to the endogenous *tsr* gene over generations under a microscope. Only cells that both were born and divided during the time-lapse were analysed. Tsr intensity at each pole was calculated with same criteria as in the EdU experiment. The accuracy of Tsr as the older pole marker was determined for each frame separately by comparing results between Tsr intensity analysis and lineage tracking. The accuracy (Fig. 4D) was shown for the last frame prior to division to mimic the EdU experiment where only cells with segregated chromosomes were analysed.

## Acknowledgements

We thank other members of the Sherratt and Uphoff group, and Katarzyna Ginda-Mäkelä for insightful discussions. The research was supported by a Wellcome Investigator Award to DJS (200782/Z/16/Z). SU was in receipt of a Henry Dale-Wellcome Fellowship.

## Author contributions

JM and DJS conceived the project and directed it. JM undertook and analyzed experiments. SU developed the microfluidics, advised on its use and was generally involved in project discussions. JM, SU and DJS wrote the paper.

## Data and code availability

All data required to understand and assess the conclusions of the research are available in the main text and supplementary materials. All materials and codes are available upon reasonable request.

## Supplementary Figures

**Fig. S1.**
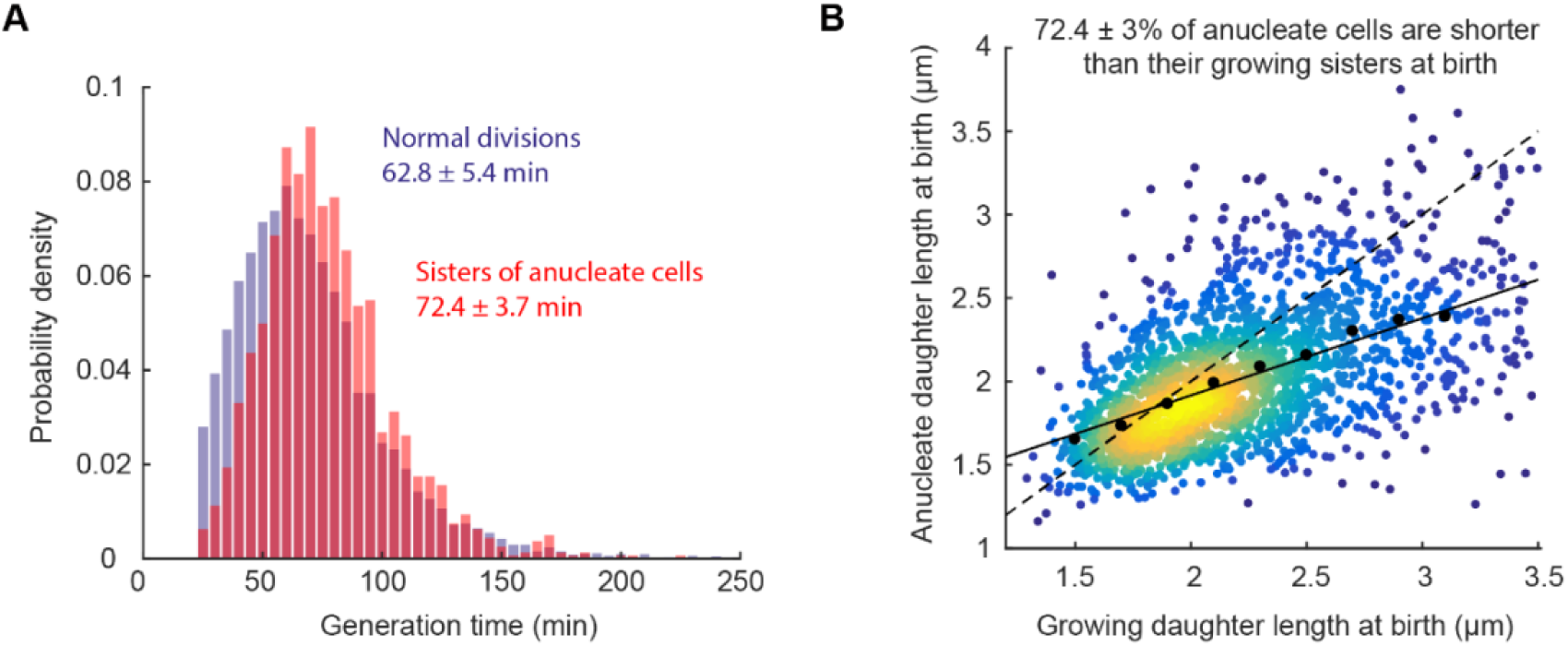
(**A**) Generation time in normally dividing cells (12103 cells) and in sisters of anucleate cells (1605 cells). Two-sample t-test of mean generation time p-value 0.2176. Data from 3 repeats. (**B**) Difference in cell length at birth between anucleate and growing sister cells at anucleate cell division. Black dashed line indicates symmetric division and solid line shows a linear fit to the data. Black circles show binned mean. 2266 cell pairs from 3 repeats.

**Fig. S2.**
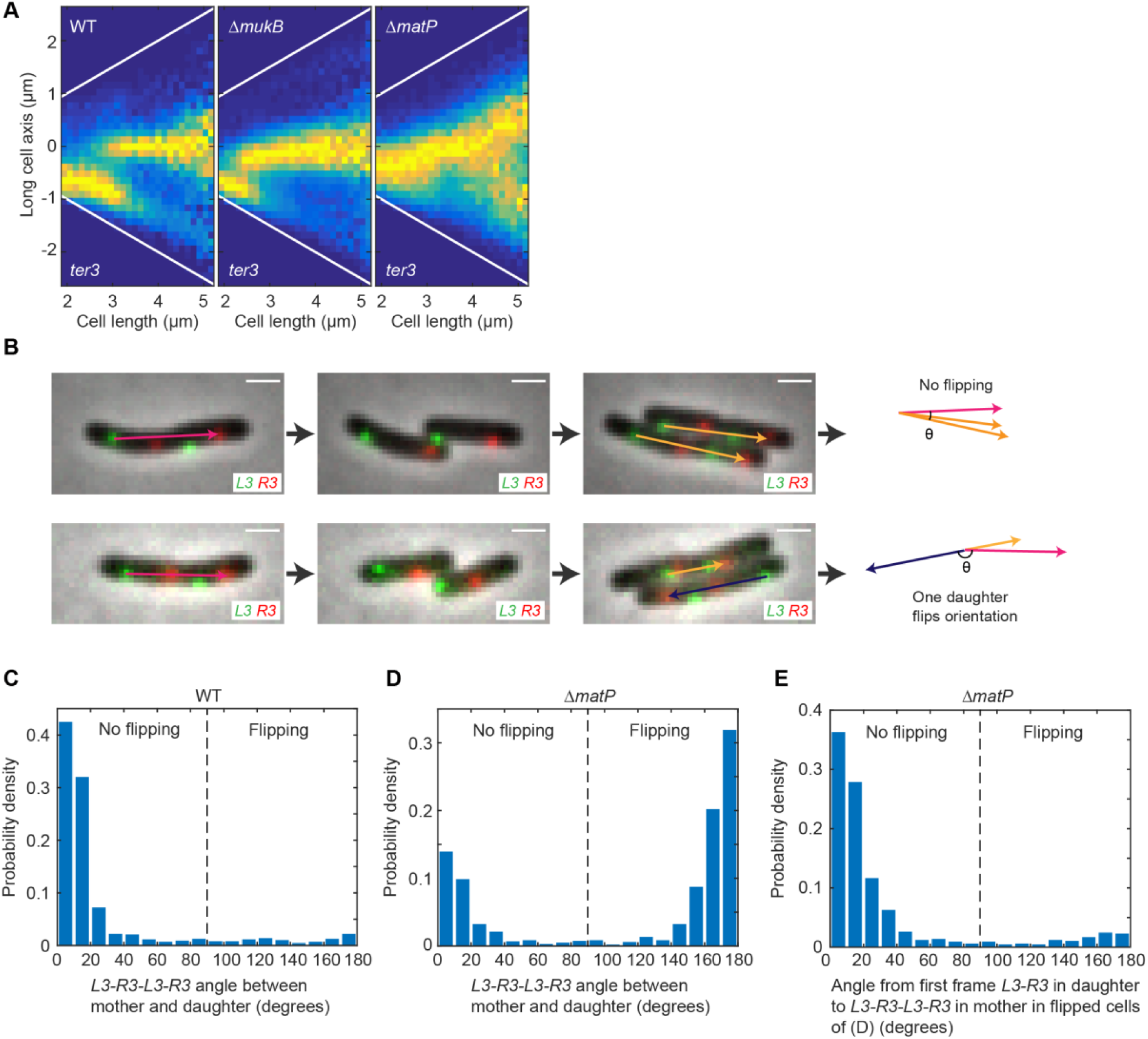
(**A**) *ter3* localization along long cell axis in WT (26926 cells), Δ*mukB* (48770 cells) and Δ*matP* (45532 cells). From same data as in Fig. 2D; *ter3* is oriented more towards the negative pole than *ori1*. Data from 3 repeats. (**B**) Representative time-lapse images of WT (top) and *ΔmatP* (bottom) cells with *L3* and *R3* markers. (top) *L3-R3-L3-R3* orientation is maintained over a generation while (bottom) *L3-R3-L3-R3* orientation is flipped. From *L3-R3-L3-R3* cells, angle between vectors pointing from the more polar *L3* to the more polar *R3* is calculated between mother cell (red arrow) and daughter cells (orange arrow) and, if the angle exceeds 90°, the chromosome orientation is considered flipped (blue arrow). Scale bars: 1 μm. Angle between mother and daughter cell *L3-R3-L3-R3* vectors in (**C**) WT (859 pairs) and (**D**) *ΔmatP* (1054 pairs) cells. Data from 3 repeats. (**E**) Angle between *L3-R3* vector in first frame of daughter cell and *L3-R3-L3-R3* vector in mother cell in flipped cells of (D). Same data is in (D).

**Fig. S3.**
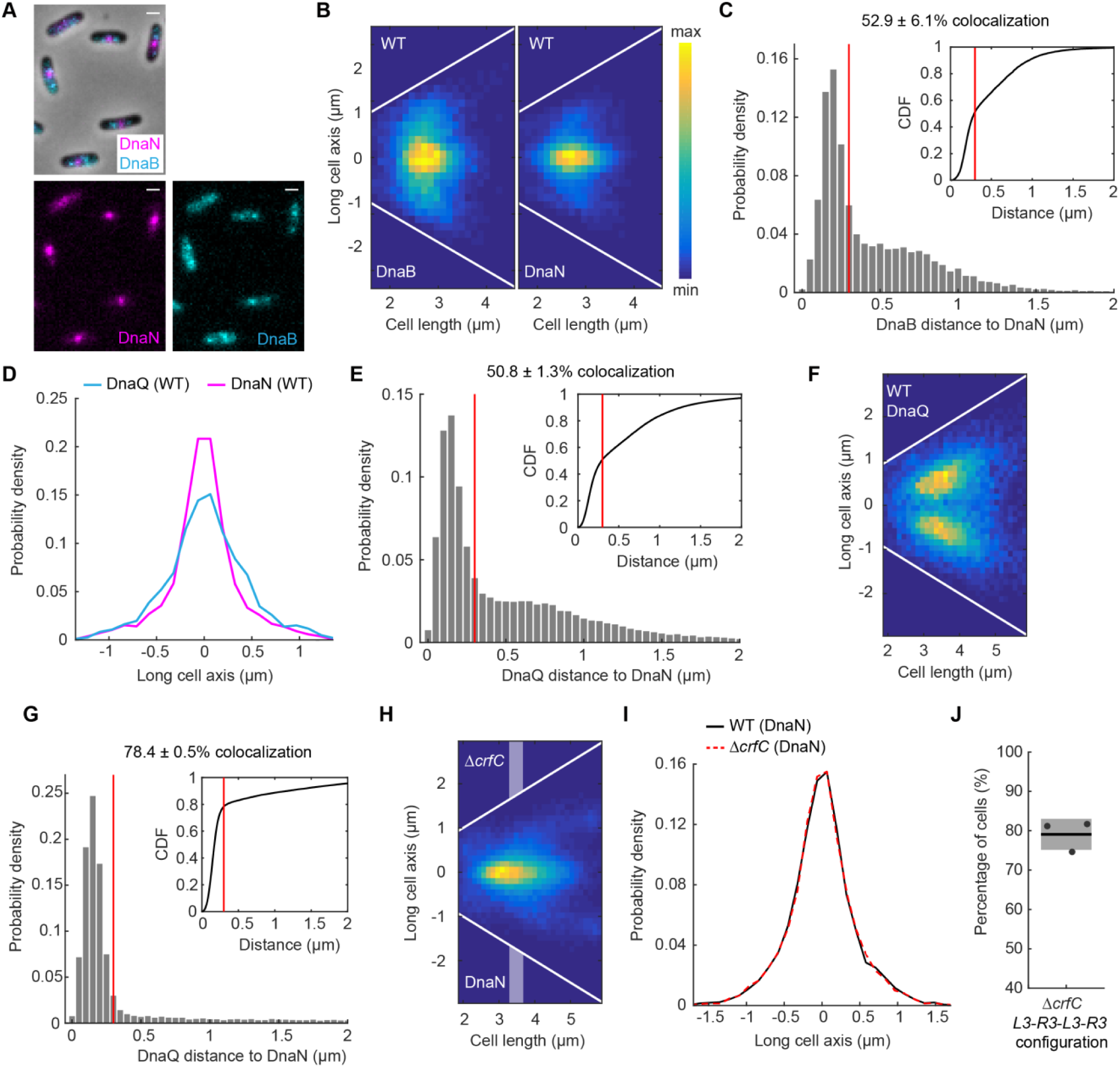
(**A**) Representative images of WT cells with labelled DnaN and DnaB. Scale bars: 1 μm. (**B**) DnaB and DnaN localization in WT cells as a function of cell length (16134 cells). White lines denote cell borders. (**C**) Distance from a DnaB locus to the closest DnaN locus. DnaB and DnaN colocalize in 52.9 ± 6.1% (±SD) of pairs (11714 pairs) as defined by a threshold (red lines) below which two proteins colocalize (dictated by a diffraction limit of 300 nm). Inset shows the same data as a cumulative distribution. Same data as in (B). (**D**) DnaQ (4567 spots) or DnaN (5393 spots) localization with early replication cells in WT (cell lengths 2.5-2.9 μm) (same data as in Fig. 3C). (**E**) Distance from a DnaQ locus to the closest DnaN locus in *ΔmatP* cells. DnaQ and DnaN colocalize in 50.8 ± 1.3% (±SD) of pairs (46330 pairs). Inset shows the same data as a cumulative distribution. Same data as in (Fig. 3F). (**F**) DnaQ localization as a function of cell length in WT cells in which DnaQ foci are spatially separate from DnaN (16158 cells). Same data as in Fig. 2C and D. (**G**) Distance from a DnaQ locus to the closest DnaN locus in *ΔmukB* cells. DnaQ and DnaN colocalize in 78.4 ± 0.5% (±SD) of pairs (32603 pairs). Inset shows the same data as a cumulative distribution. Same data as in (Fig. 3G). (**H**) DnaN localization in *ΔcrfC* cells as a function of cell length (49955 cells). Shaded areas denote intermediate cell lengths for localization data in (I). White lines denote cell borders. (**I**) DnaN localization with intermediate cell lengths (3.3-3.7 μm) in WT (8006 spots) and *ΔcrfC* (10691). Data from (H) and Fig. 3E. (**J**) Percentage of *ΔcrfC* cells (8393 cells) with *L3-R3-L3-R3* (or *R3-L3-R3-L3*) configuration (versus *L3-R3-R3-L3* or *R3-L3-L3-R3*) in double *L3* and *R3* focus cells. All data from 3 repeats.

**Fig. S4.**
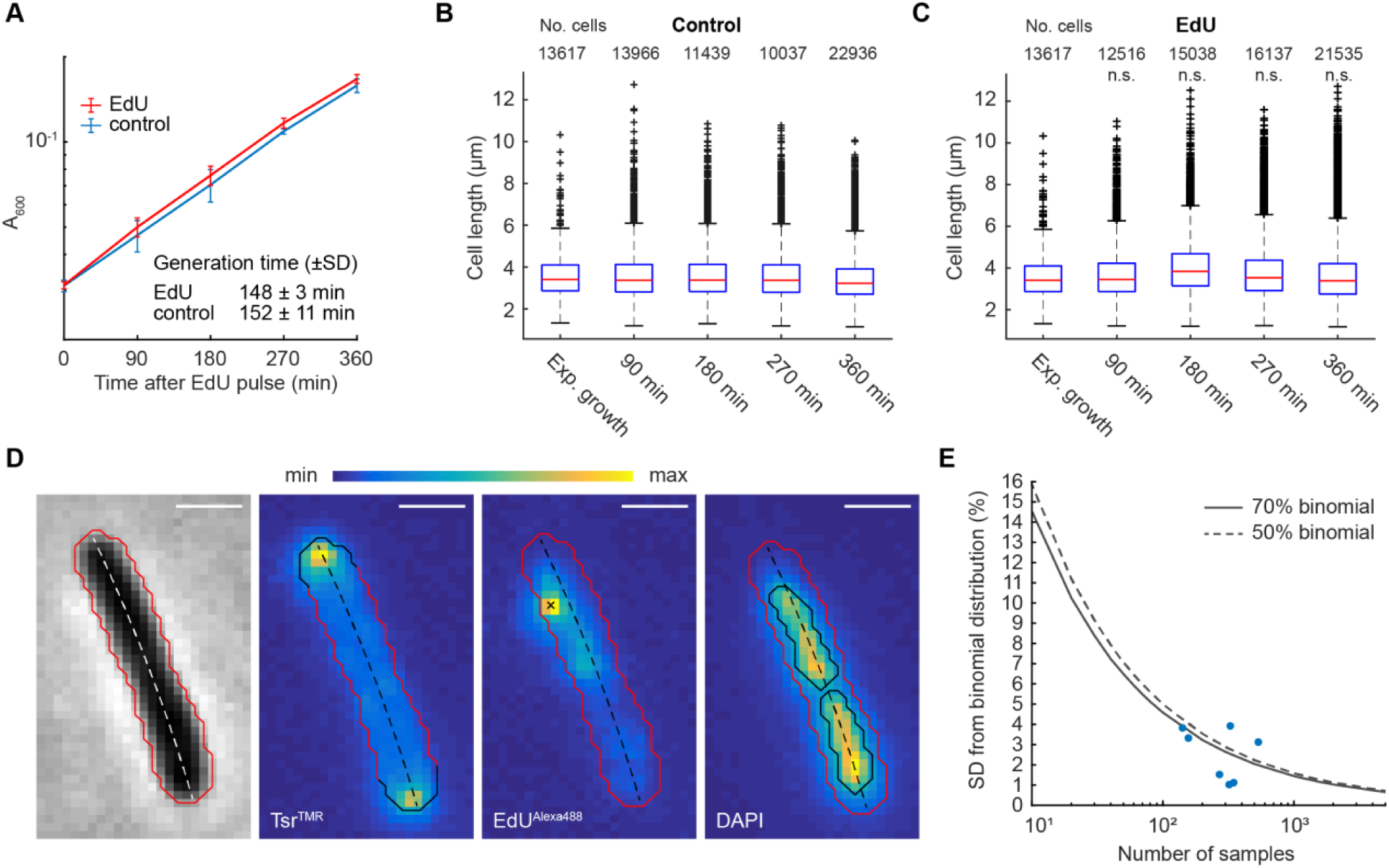
(**A**) Cell growth following a 15 min EdU pulse compared to no EdU. Cell length at different time intervals (**B**) without or (**C**) with EdU pulse. n.s. indicates two-sample t-test of mean cell length compared to control p-value > 0.01. Data from 3 repeats. (**D**) Image analysis of EdU experiment. A representative cell after EdU protocol showing Tsr^TMR^, EdU^Alexa488^ and DAPI labelling. Red line is the cell border and dashed line shows the center line of the cell. Black line in Tsr^TMR^ channel shows the pole areas from where the Tsr intensity is calculated. Black cross in EdU^Alexa488^ channel indicates a detected EdU focus. Black lines in DAPI channel indicate segmented nucleoid areas. For more information see Methods. Scale bars: 1 μm (**E**) Accuracy of the retention measurement as function of sample size. Different sample sizes were drawn from a binomial distribution with 50% (dashed line) or 70% (solid line) success rate and SD was calculated between them (10^5^ repeats for each value). The data from Fig. 5 are shown with dots.

**Table S1.**
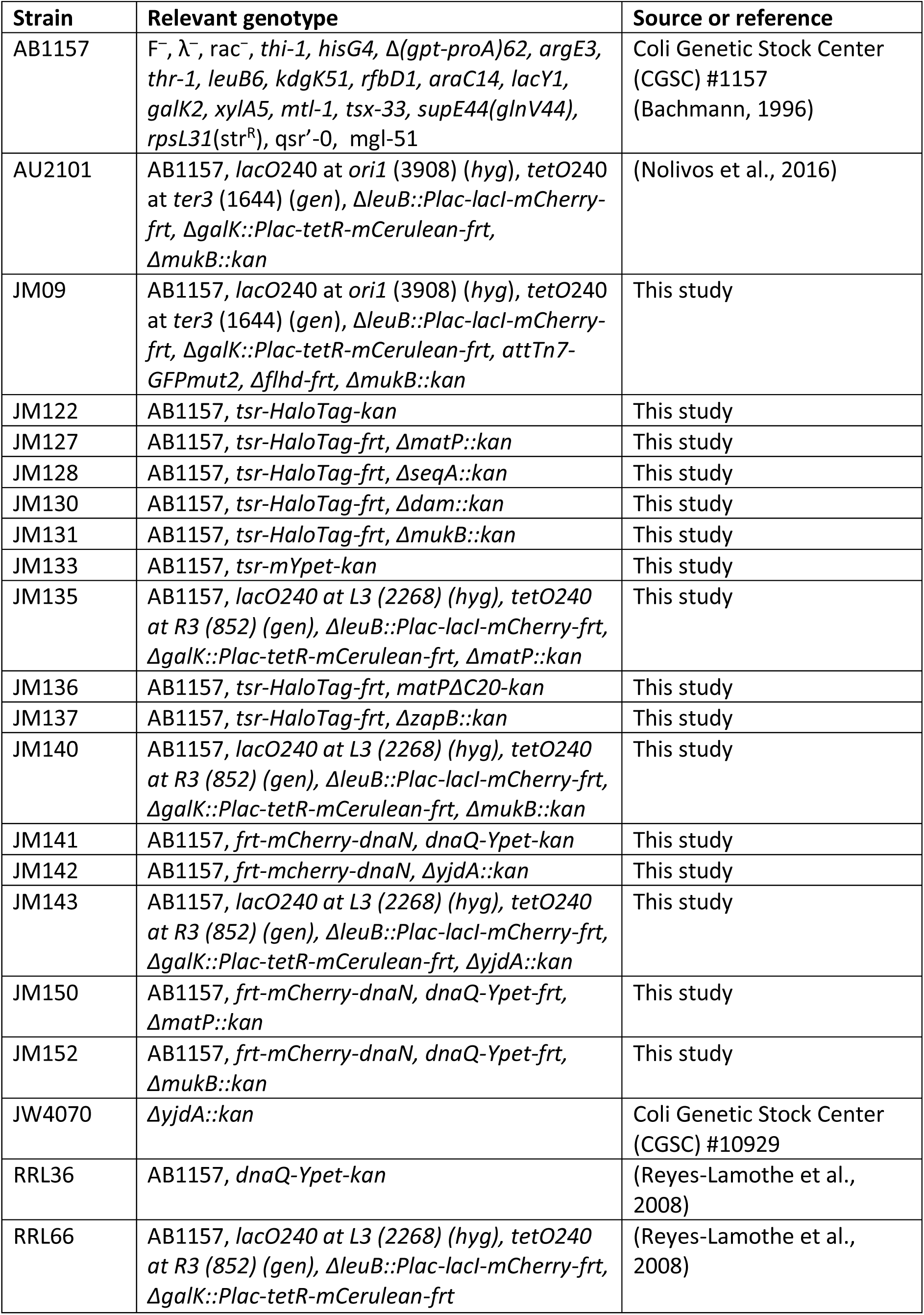
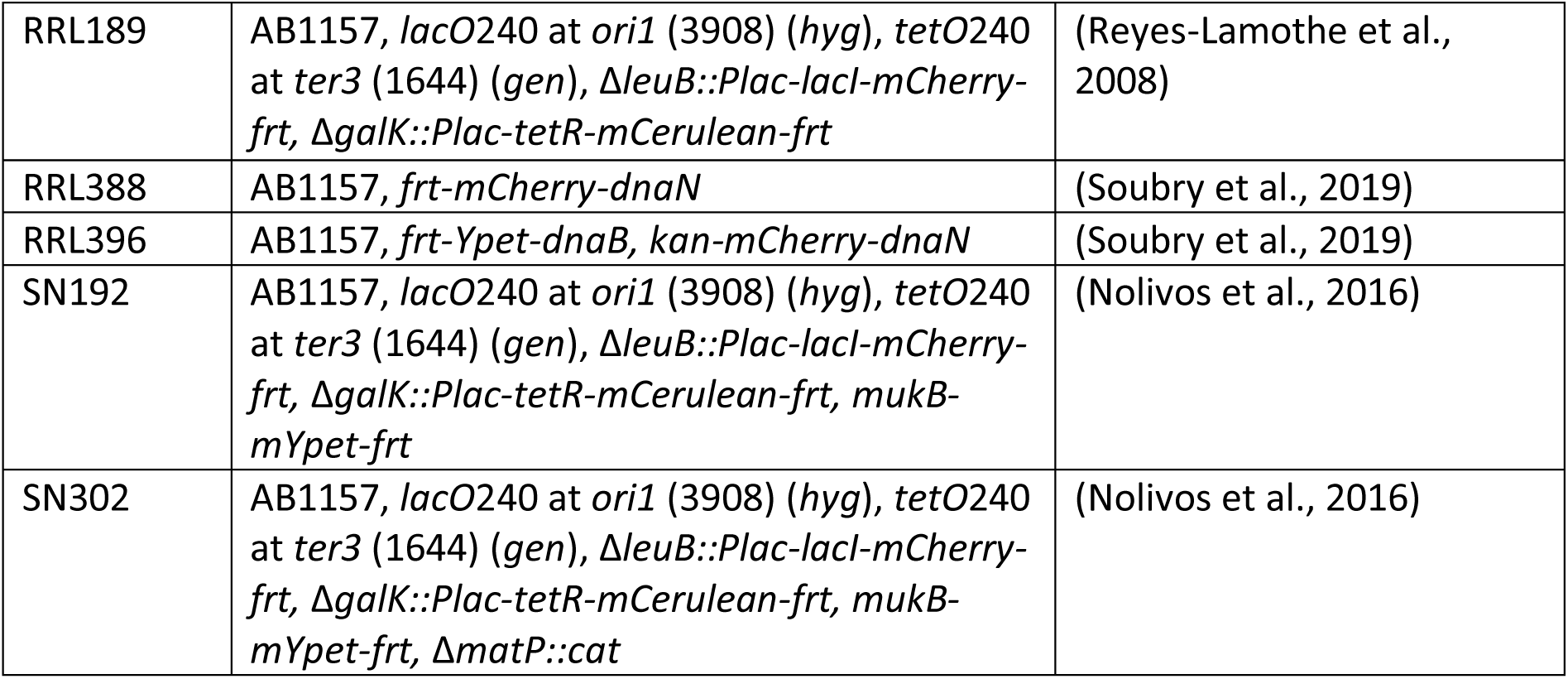
Strain list.

**Table S2.**
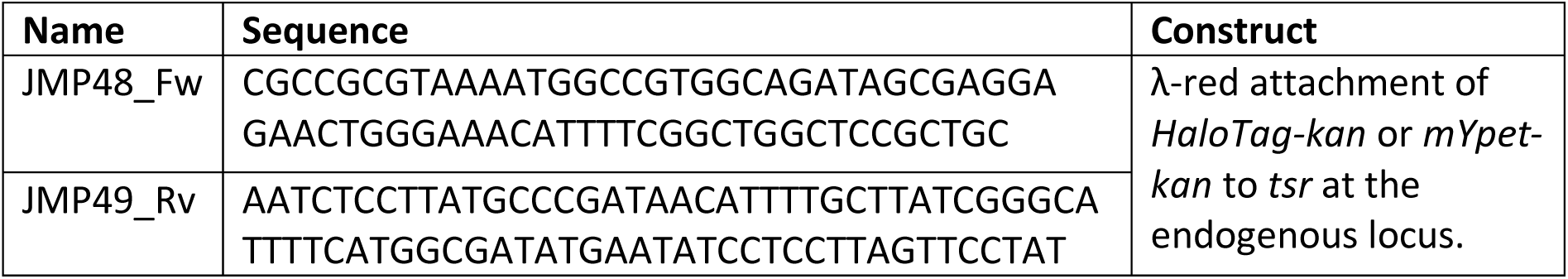
Primer list

